# NMFClustering: Accessible NMF-based clustering utilizing GPU acceleration

**DOI:** 10.1101/2023.06.16.545370

**Authors:** Ted Liefeld, Edwin Huang, Alexander T. Wenzel, Kenneth Yoshimoto, Ashwyn K Sharma, Jason K Sicklick, Jill P Mesirov, Michael Reich

## Abstract

**Summary:** Non-negative Matrix Factorization (NMF) is an algorithm that can reduce high dimensional datasets of tens of thousands of genes to a handful of metagenes which are biologically easier to interpret. Application of NMF on gene expression data has been limited by its computationally intensive nature, which hinders its use on large datasets such as single-cell RNA sequencing (scRNA-seq) count matrices. We have implemented NMF based clustering to run on high performance GPU compute nodes using CuPy, a GPU backed python library, and the Message Passing Interface (MPI). This reduces the computation time by up to three orders of magnitude and makes the NMF Clustering analysis of large RNA-Seq and scRNA-seq datasets practical. We have made the method freely available through the GenePattern gateway, which provides free public access to hundreds of tools for the analysis and visualization of multiple ‘omic data types. Its web-based interface gives easy access to these tools and allows the creation of multi-step analysis pipelines on high performance computing (HPC) clusters that enable reproducible *in silico* research for non-programmers.

**Availability and Implementation:** NMFClustering is freely available on the public GenePattern server at https://genepattern.ucsd.edu. Code for the NMFClustering is available under a BSD style license on github at https://github.com/genepattern/nmf-gpu.

**Contact:** Ted Liefeld

**Supplementary Information:** Supplementary data are available at Bioinformatics online and at https://datasets.genepattern.org/?prefix=data/test_data/NMFClustering/.

## Background

Non-negative matrix factorization (NMF) (Lee and Seung 2001) has proved to be a very effective technique for decomposing array-based biological data into meaningful components. In gene expression analysis, NMF is used to describe the activity of tens of thousands of genes in a typical gene expression dataset in terms of a small number, *k*, of metagenes, i.e., positive linear combinations of genes (Brunet et al. 2004). Biological samples data can then be summarized as expression patterns of their metagenes. Compared with other dimension reduction techniques such as Principal Components Analysis (PCA), the components identified by NMF are more easily related to actual biological processes and mechanisms due to the non-negativity constraint.

However, the use of NMF has been limited by its computational requirements. NMF is a non-convex optimization problem that requires many iterations to converge to a solution, and the solution is not guaranteed to be a global minimum. Compounding this problem is that the optimal number of clusters, k, is seldom known in advance. Therefore, when NMF is used to cluster data, it is run many times with varying initial conditions over a range of values of k. Cluster stability metrics such as the cophenetic correlation coefficient (Farris 1969) or the silhouette score (Rousseeuw 1987) are then used to identify the value of k that corresponds to the most stable clustering.

The result is that on top of the computational complexity of NMF, to perform clustering it must be run many thousands of times for a given analysis. With the current NMF-based methods it is not unusual to encounter analyses that require days to run on a dataset as small as 1000 cells by 5000 transcripts. For larger gene expression or single-cell RNA-Seq (scRNASeq) datasets, use of NMF becomes impractical, requiring potentially weeks of computing time on large-memory supercomputers.

Graphical Processing Units (GPUs) have become a popular and cost-effective way to speed up computation. GPUs are limited in memory compared to CPUs but are designed for processing large blocks of data in parallel and are particularly well-suited to matrix operations, which are the basis of the NMF algorithm. Via GPU application programming interfaces (APIs) such as the nVidia Compute Unified Device Architecture (CUDA)(NVIDIA 2007), general purpose algorithms can be executed in parallel using GPUs to speed computation. To further parallelize computation across multiple GPUs, parallel programs can be synchronized using the Message Passing Interface (MPI).

Mejía-Roa (Mejia-Roa et al. 2015) demonstrated NMF-mGPU, a C-language implementation of NMF that used CUDA and MPI to perform multi-GPU computation of NMF. This implementation allowed processing up to 120 times faster than conventional single-CPU implementations and demonstrated the capability of processing matrices up to 54000 × 2000 in size. This implementation included the ability to split the NMF computation across multiple GPUs to permit the calculation to be performed on datasets larger than could be held in GPU memory. However, this program was not easily usable by many biomedical sciences researchers as it required specialized programming skills and access to an HPC cluster..

GenePattern (Reich et al. 2006), www.genepattern.org, is a gateway providing free access to cloud-based and HPC systems. GenePattern includes access to hundreds of analysis tools (modules) and visualization tools for multiple genomic data types without requiring any programming skills. It has a web-based interface to provide easy access to these tools and allows the creation of multi-step reproducible *in silico* research workflows. These can be implemented as GenePattern analysis pipelines, or alternatively as Jupyter (Ragan-Kelley et al. 2014) Notebooks via the GenePattern Notebook Environment (Reich et al. 2017). The publicly available GenePattern servers currently support thousands of users and log up to tens of thousands of analyses each month which are executed on clusters in the AWS Cloud or on the Expanse (Strande et al. 2021) supercomputer cluster at the UC San Diego Supercomputing Center (SDSC). GenePattern has long included an R-language implementation of the NMF Clustering algorithm as described in Brunet et al. as the NMFConsensus GenePattern module. This implementation served as a baseline for performance comparison.

To bring the power of the Meija-Roja NMF-GPU implementation to non-programming investigators, we have reimplemented their algorithm using CuPy (Okuta et al. 2017) an open-source library for GPU-accelerated computing with Python. It is freely available on GenePattern as the NMFClustering module at https://genepattern.ucsd.edu and is also distributed as both open source code and as a Docker (Boettiger 2015) container. For HPC clusters, the Docker container can be cross-compiled into Singularity (Kurtzer, et al. 2017) if required. By wrapping this implementation in GenePattern, we have made it available to all investigators without the need for any programming skills and have also provided access to GPU cycles for the processing of their data.

## Results

We illustrate the improved performance of NMFClustering, our CuPy based NMF-GPU implementation, relative to the previous generation of NMFConsensus implemented in the R programming language by demonstrating its ability to handle much larger datasets, and by demonstrating its improved performance on smaller datasets. It is not possible to do a true performance comparison on larger datasets as the R language NMFConsensus implementation requires in excess of 2 days, the time limit for GPU jobs on the Expanse HPC cluster, to analyze datasets larger than 1,000 cells/samples. We note that most single cell data sets are much larger than this.

For our test dataset, we use a single-cell RNA-Seq dataset derived from patient samples of gastrointestinal stromal tumors (GIST). The complete dataset is a count matrix of 77020 cells and 28561 transcripts. To create datasets small enough to run on the R-language NMFConsensus, we subsetted this dataset, creating smaller versions with 5,000 transcripts and cell counts between 20 and 1,000. Additional larger datasets with 8,000 to 32,000 cells were run through NMFClustering in a CUDA configuration using a single node and both single and 2 GPUs allocated. Finally two larger datasets of 64,000, and 77,020 (the complete dataset) cells were created and run using a CUDA configuration of 4 nodes and 4 GPUs.

Using these datasets we ran the NMFConsensus module on GenePattern’s public cloud based server, which launched AWS C5 compute nodes of various types through AWS Batch using the SPOT_CAPACITY_OPTIMIZED strategy. Datasets larger than 1,000 cells were not run using the CPU based algorithm as they were unable to complete the analysis below the 48 hour time limit for most jobs on the SDSC Expanse system. The Expanse GPU nodes have dual 20 core Intel Xeon processors with 384 GB of memory and four NVIDIA V100 processors (32 GB SMX2). Each dataset size was run 4 times and the average elapsed time used for the following analysis.

For the NMFClustering code we ran the same datasets as were used for NMFConsensus using the GPU nodes of the SDSC Expanse supercomputer. Using a single GPU configuration we ran up to 8,000 cells successfully. The largest datasets consistently failed with exhausted GPU memory for the single GPU configuration. Since the NMFClustering implementation could split the dataset to compute only a portion on each GPU at one time, using MPI to coordinate, we could reduce the impact of limited GPU memory by adding GPUs to the runtime configuration. Therefore for the largest 4 datasets (16,000 through 77,020 cells) we ran using a quad-GPU (4 GPU) configuration which successfully completed a NMF-GPU run in under 7 hours on the largest dataset. The dataset size and runtimes, averaged over 4 runs, are displayed below in figure 1.

**Figure 1.**
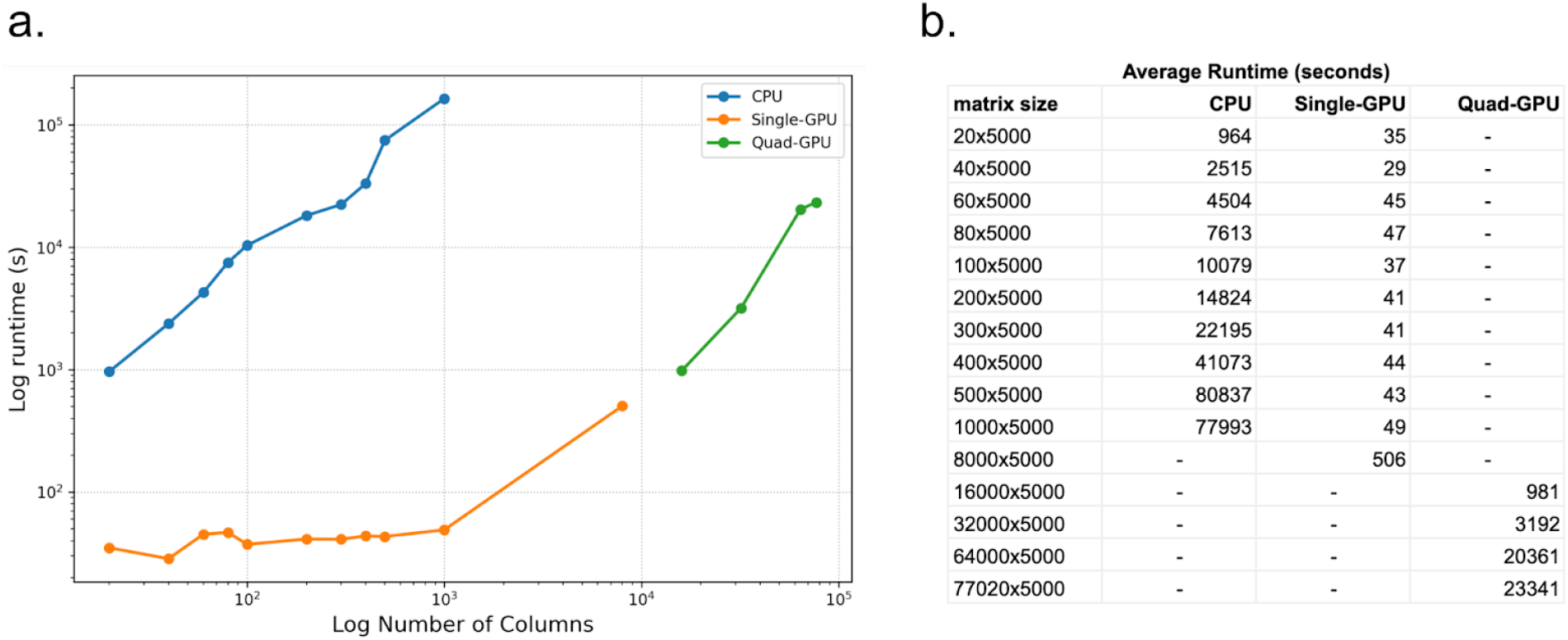
Plot (a) and table (b) of compute time vs number of cells (columns) for CPU-based NMFConsensus and multiple GPU configurations of NMFClustering. Each dataset contains 5000 rows and was run 4 times to ensure consistency of runtimes.

With 1000 cells (columns) and 5000 transcripts (rows), the largest dataset that could be computed on the CPU, the average execution time was 77993 seconds. The same dataset completed using NMFClustering on a single GPU in 49 seconds, 1588 times faster or in 0.062% of the time of the CPU version. The smallest improvement was for the smallest 20 sample dataset, where the GPU speedup was 27 times faster than the CPU version.

The NMFClustering (GPU-based) and NMFConsensus (R-based) implementations are available as modules on the public GenePattern server at https://genepattern.ucsd.edu. The R code for the NMFConsensus GenePattern module is available in github at https://github.com/genepattern/NMFConsensus. The Python code for the NMFClustering (GPU based) implementation is also available on github at https://github.com/genepattern/nmf-gpu.

## Features

The NMFClustering module, and underlying Python code, allow the user to define the minimum and maximum values of k, the number of clusters. It also allows the user to select the number of NMF runs with varied initial conditions per value of k, called the “num clusterings” parameter as well as the maximum number of total iterations of NMF to run per clustering to prevent the process for running forever if it does not converge to a stable result. To control the convergence it provides parameters for the “stop convergence”, to specify how many “no change” checks are needed to stop NMF iterations before max iterations is reached along with the “stop frequency” parameter, which defines the frequency (number of NMF iterations) between ‘no-change’ checks. Finally the module also allows the user to define the maximum error difference for KL divergence.

For advanced users, the module allows the selection from three parallelization strategies. They are: (1) ‘serial’ in which no parallelization is used; (2) ‘kfactor’ where NMF runs on values of k are balanced sending the entirety of runs for one value of k to a single GPU; and (3) ‘Input matrix’ parallelization in which case the input matrix is broken up with each GPU computing on only a portion of the input as defined in Meija-Roja et al, 2015. The ‘input matrix’ serialization is required for datasets that are too large to fit in the relatively smaller memory space of the GPUs and is what we used for the larger datasets in the performance testing.

## Outputs

For each value of k, NMFClustering outputs a consensus matrix, which contains the number of times each sample appears in the same cluster as each other sample. In a stable clustering, the same samples will cluster with each other a majority of the time. Consensus matrices are output in the GenePattern gct[14] tab delimited format. If the number of input columns is under 1,000 the module also provides sorted consensus matrices in gct format, and if less than 100 columns, the consensus matrices will be plotted and returned as a pdf file.

## Documentation

Documentation for NMFClustering is available on github pages at https://genepattern.github.io/NMFClustering/v3/.

## Acknowledgements

The work was supported by the National Institutes of Health U24CA194107 to JPM, R50CA243876 to TL, F31CA257344 to ATW, R01CA226803 to JKS, and T32CA121938 fellowship to AKS.

This work used the Expanse supercomputer at the San Diego Supercomputer Center through allocation MED220012 from the Advanced Cyberinfrastructure Coordination Ecosystem: Services & Support (ACCESS) program, which is supported by National Science Foundation (grant numbers #2138259, #2138286, #2138307, #2137603, and #2138296).

## Supplemental Data

### NMFClustering User Interface

**Figure S1.**
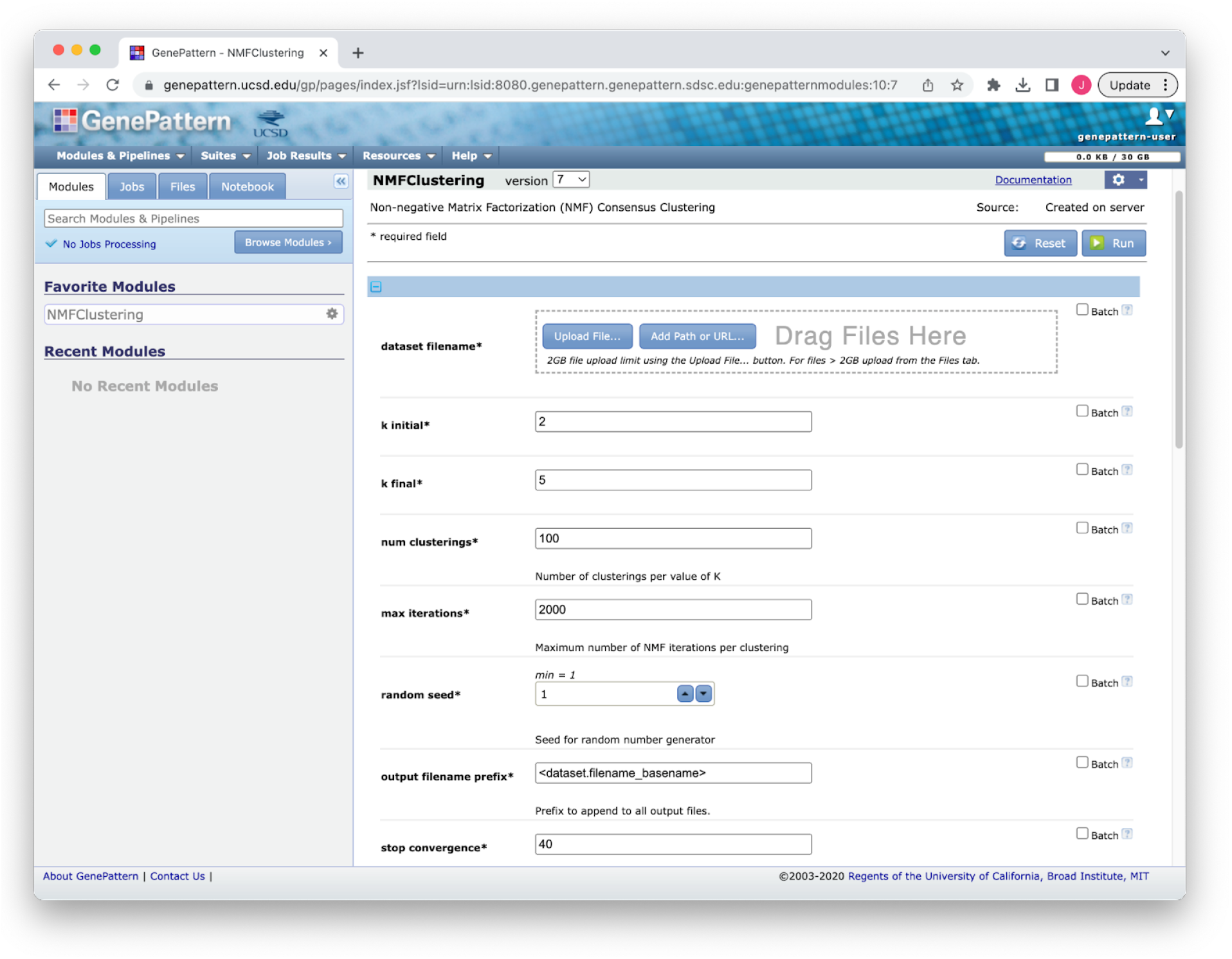
NMFClustering interface in GenePattern showing a subset of the available parameters

### Example Consensus Matrix plot

**Figure S2.**
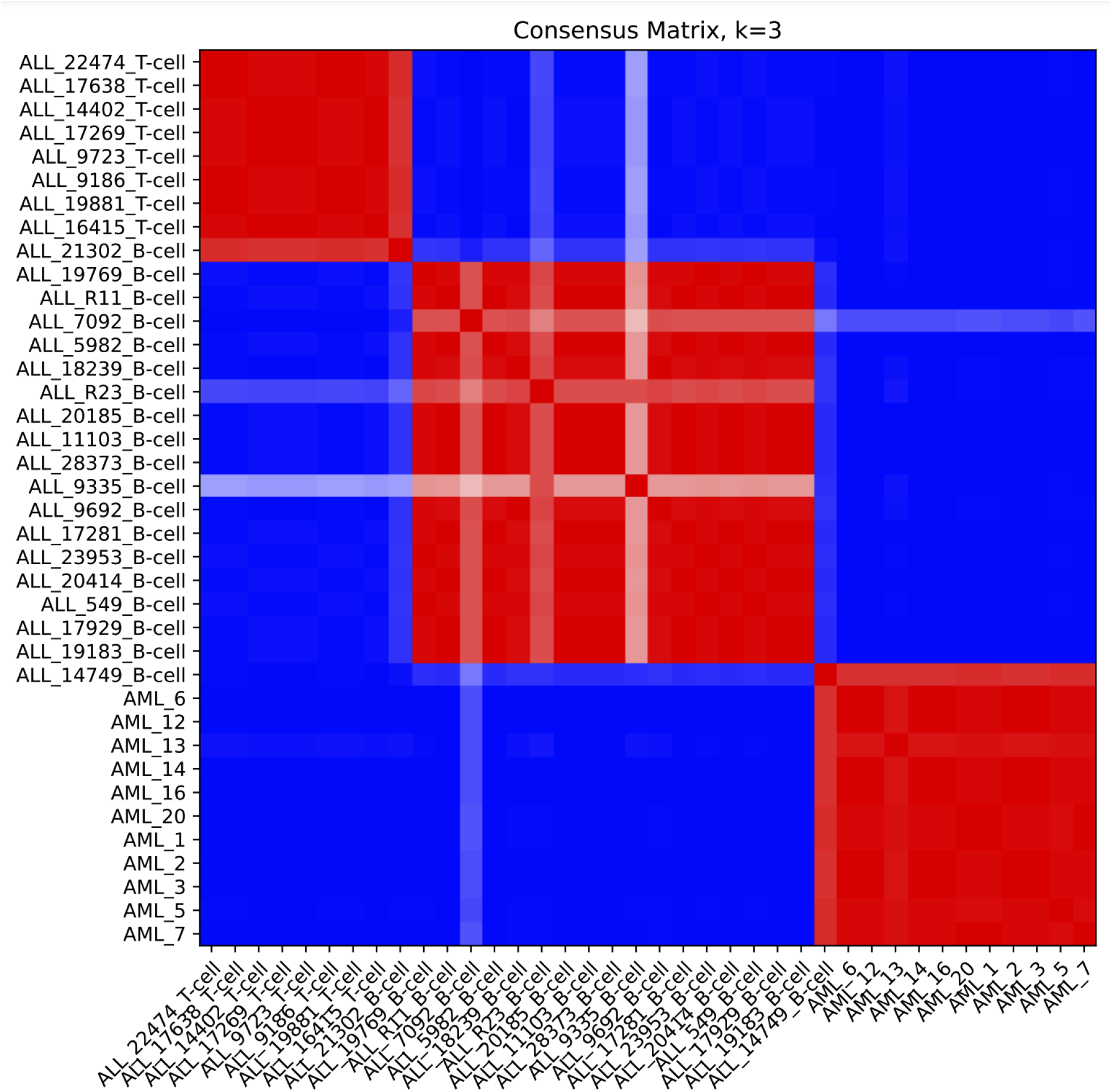
Example consensus matrix plot for k=3 using ALL_AML_data.gct dataset from the sample test data folder (below). Compare to figure 4, panel a in Brunet et al.

### Sample Test Data

Test datasets are available at https://datasets.genepattern.org/?prefix=data/test_data/NMFClustering/

## Notes

https://datasets.genepattern.org/?prefix=data/test_data/NMFClustering/

### Competing Interest Statement

The authors have declared no competing interest.

### Summary of Updates

Updated the acknowledgements section, slight reformatting and added additional grants.

